# Worldwide population genomics reveal long-term stability of the mitochondrial chromosome composition in a keystone marine plant

**DOI:** 10.1101/2023.04.21.537793

**Authors:** Marina Khachaturyan, Thorsten B. H. Reusch, Tal Dagan

## Abstract

Mitochondrial genomes (mitogenomes) of flowering plants are comprised of multiple chromosomes. Their copy number and composition can be dynamic within and among individual plants due to uneven replication of the chromosomes and homologous recombination. Nonetheless, despite their functional importance, the level of mitogenome conservation within species remains understudied. Whether the ontogenetic variation translates to evolution of mitogenome composition over generations is currently unknown. Here we show that the mitochondrial chromosome composition of the seagrass *Zostera marina* is conserved among worldwide populations that diverged ca 350,000 years ago. Using long-read sequencing we characterized the *Z. marina* mitochondrial genome and inferred the repertoire of recombination-induced configurations of its eight chromosomes. To characterize the chromosome composition worldwide and study its evolution we examined the mitogenome in *Z. marina* meristematic region sampled in 16 populations from the Pacific and Atlantic oceans. Our results reveal a striking similarity in the chromosome copy number suggesting stable equal proportions among distantly related populations and a high conservation of the mitochondrial genome within the plant germline, despite a notable variability during individual ontogenesis. Our study supplies a link between observations of dynamic mitogenomes at the level of plant individuals and long-term mitochondrial evolution.

**Significance statement:** Extensive studies on evolution of plant mitochondria in individual plants revealed great variability of the mitogenome architecture across tissues, however, data on the mitochondrion evolution at the population level is still scarce. We show that the mitochondrial genome architecture in a keystone marine plant, *Zostera marina*, remained conserved over ca. 350,000 years worldwide. We suggest that the extreme conservation of the *Z. marina* mitochondria is a manifestation of streamlined mitochondria inheritance over plant generations, e.g., via a plant germline.

## Introduction

Plant mitochondrial genomes are often described as a single circular DNA molecule due to their homology to bacterial chromosomes and common convention for the description of most animal mitogenomes. However, a number of recent experiments questioned this dogma revealing a variety of multipartite mitogenomes comprising several DNA molecules (reviewed in Sloan 2013 and Wu et al. 2020). Mitochondrial chromosomes can be circular, linear, or branched, where a single mitogenome can be composed of DNA molecules that vary in shape, size, and functionality (Sloan 2013; Arimura 2018; Kozik et al. 2019). Most complete mitogenomes comprise up to 5 chromosomes (Wu et al. 2020), yet up to 63 chromosomes were reported in the terrestrial eudicot *Silene noctiflora* (Wu et al. 2015). Plant mitochondrial chromosomes often contain repeated regions, which serve as homologous recombination sites. Repeats of intermediate size (e.g., 50-500bp) have low recombinational activity, whilst large repeats are highly conducive to recombination events, hence are considered recombinationally active (Brieba 2019; Sullivan et al. 2020). Chromosomes containing such recombinationally active repeats are therefore prone for recombination events that change their conformation. For example, events of reciprocal recombination via a pair of direct recombinationally active repeats in a circular chromosome, termed the master circle, lead to the formation of two smaller DNA molecules, termed subgenomes, that are considered as mitochondrial chromosomes as well (first described in Palmer and Shields 1984; reviewed in Sloan 2013 and Gualberto and Newton 2017). In some plants the multipartite mitogenome is formed solely by a single master circle and a set of corresponding subgenomes (Alverson et al. 2011; Guo et al. 2017; Dong et al. 2018). In several plant species the mitogenome includes multiple master circles and the corresponding subgenomes (Sloan et al. 2012; Shearman et al. 2016; Shtratnikova et al. 2020). Linear mitochondrial chromosomes may include telomere-like ends, that are visible in the mitogenome assembly (Shtratnikova et al. 2020). At the same time, chromosomes assembled as a circular configuration may correspond to a combination of overlapping linear forms *in vivo*, which assist a recombination-dependent replication of the mitogenome (Gualberto and Newton 2017; Arimura 2018).

Notwithstanding above insights, mitochondrial chromosomes still are traditionally described as circular, although the predominance of circular configurations was proposed only for meristematic tissues (Arrieta-Montiel et al. 2001; Preuten et al. 2010; Woloszynska 2010; Mower, Case, et al. 2012), which are distinct from other tissues due to their contribution to the plant germline (Kwiatkowska 2008; Burian 2021). The mitochondrial main genome is thus composed of chromosomes that correspond to either master circles, subgenomes or linear chromosomes. In addition, occasional recombination events may lead to the formation of a dynamic variable reservoir of substochiometric forms, which are alternative conformations that persist in a low copy number (Small et al. 1987; Alverson et al. 2011).

Plant mitochondria constantly undergo fusion and fission that facilitate genome mixing among mitochondria within the cell (reviewed in Rose 2019 and Rose 2021). The genome intermingling may lead to individual mitochondria having an incomplete set of chromosomes (Preuten et al. 2010), therefore any mitogenome should be considered at the cellular rather than the organellar level (Lonsdale et al. 1988; Rose 2021). Another notable characteristic of mitochondrial DNA is that its replication is not coordinated with cell division (termed relaxed replication; Birky, Jr. 1994). One implication of relaxed replication is the possibility for an uneven replication of different mitochondrial chromosomes, which may change chromosome stoichiometry following cell division (Woloszynska 2010). Such changes in the relative abundance of the main genome chromosomes are termed stoichiometric shifting (Gualberto and Newton 2017). In contrast, transitions between the main genome and substoichiometric forms are termed substoichiometric shifting; these occur following a dramatic increase in the relative copy number of a specific substoichiometric form, or a dramatic decrease in the relative copy number of a main genome chromosome (Small et al. 1987; reviewed in Mackenzie 2007). Notably, chromosomes stoichiometry and its dynamics remains elusive for most thoroughly constructed multipartite mitogenomes through evolutionary time, e.g. among populations of plant species.

Previous studies suggest that mitochondrial genetic variation due to (sub)stoichiometric shifting may correspond to somatic variation within individual plants (Suzuki et al. 1996; Bartoszewski et al. 2004; Sun et al. 2012; Woloszynska et al. 2012; Sloan 2013). The dynamics of recombination events and uneven replication of the mitochondrial chromosomes have the potential to manifest in (sub)stoichiometric shifting. To the best of our knowledge, only a single study of (sub)stoichiometric shifting at the population level was performed thus far, focusing on *S. noctiflora*; the study revealed signatures of ongoing mitogenome reduction that manifests in chromosome loss (Wu and Sloan 2019). Nonetheless, the evolution of *S. noctiflora* mitogenome may represent an extreme case of a large and highly fragmented genome; since some chromosomes in *S. noctiflora* carry no genes, genome reduction due to substoichiometric shifting may be neutral. The effect of (sub)stoichiometric shifting on the evolution of plant mitogenomes thus remains understudied and requires observations of mitogenome diversity across populations.

Here, we study the mitochondrial chromosome stoichiometry of the marine plant *Zostera marina* (eelgrass). This species served recently as the focus of investigations on monocotyledonous adaptation to marine habitats and species radiation in the ocean (Olsen et al. 2016; Ma et al. 2021; Yu et al. 2022). *Z. marina* reproduces both sexually and via vegetative (clonal) reproduction. The latter leads to the formation of large eelgrass clones that can be several hectares in extent (Yu et al. 2020). A recent study of *Z. marina* samples collected in 16 locations worldwide revealed that all extant populations originated in the North-West Pacific and started colonizing Pacific and Atlantic oceans ca. 350kya (Yu et al. 2022). The availability of high-resolution genomic information at the population level including a dated phylogeny enabled us to study the evolutionary dynamics of mitogenome composition of a keystone marine plant.

## Results

### *Zostera marina* mitochondrial genome assembly reveals two main chromosomes

The mitogenome sequence of *Z. marina* was previously reported to include a single chromosome (Petersen et al. 2017). The availability of PacBio long-read sequence data that was recently used to assemble the *Z. marina* nuclear genomes to chromosmal resolution (Ma et al. 2021) enabled us to also assemble the mtDNA with high accuracy. Mitochondria contigs were identified by sequence similarity to the published mitogenome and used for genome assembly. The presence of ambiguous edges in the assembly graph suggested a complex mitogenome structure with several chromosomes. The assembly graph consisted of three edges of 149,304 bp, 35,405 bp, and 2,339 bp lengths (Figure 1A). As the latter edges are alternative assembly solutions, it was impossible to reconstruct the genome as a single chromosome unless a duplication of 149,304 bp is proposed. As a more parsimonious solution, we therefore resolved the mitochondrion genome into two circular chromosomes Chr1 and Chr2 of 184,709 bp and 156,143 bp, respectively. The alternative edges are unique parts of the chromosomes and the largest edge is the common part. Since no variant sites were found in the short read data of the parts common to both chromosomes in the assembly, we concluded that these parts are identical. Our assembly of the *Zostera marina* mitogenome thus extends the set of known multichromosomal plant mitogenomes that possess a unique configuration composition.

**Fig. 1:**
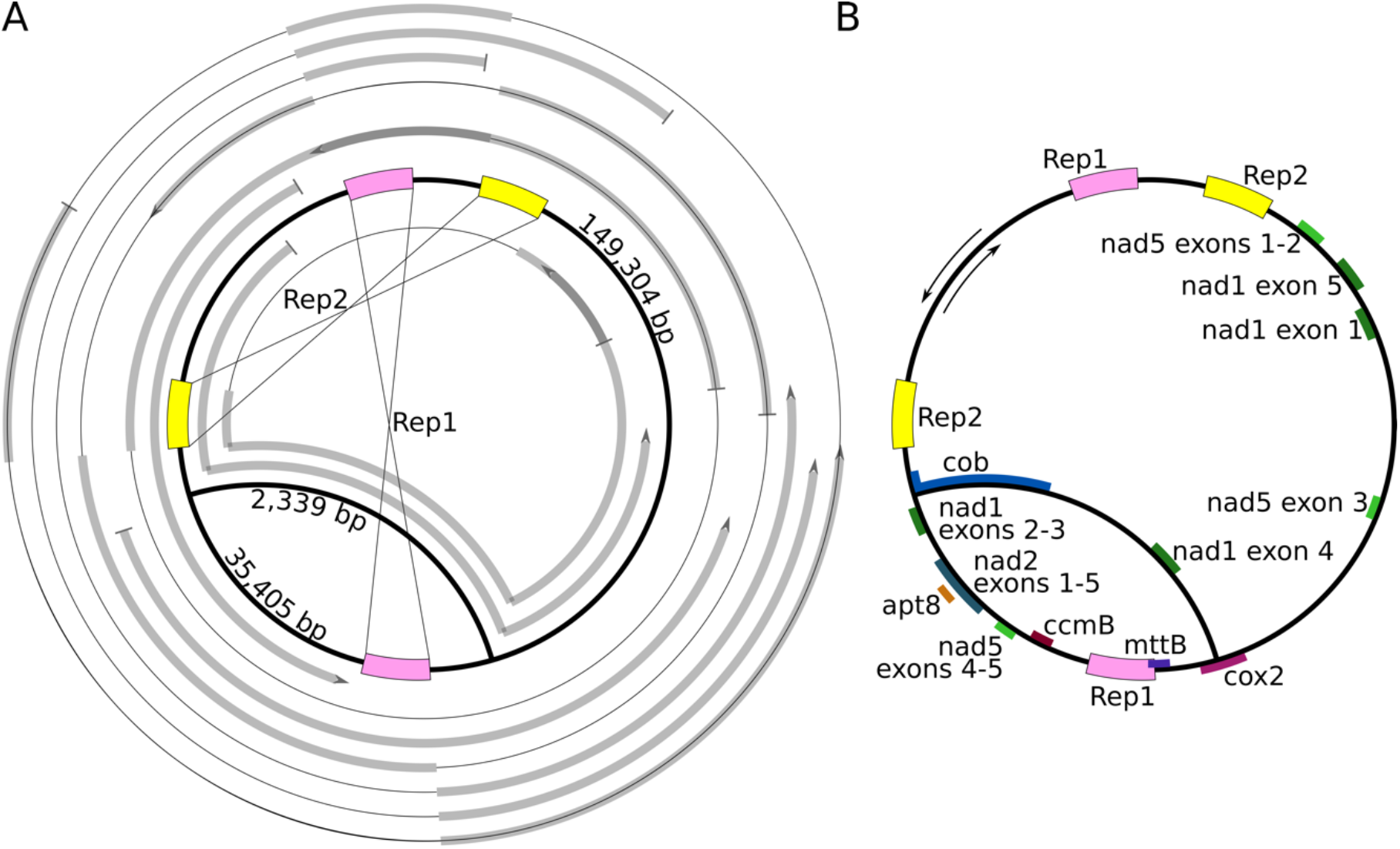
*Z. marina* mitochondrion genome assembly and annotation. a, The nine mitochondrial PacBio contigs aligned to the proposed mitochondrion assembly graph. The assembly graph is depicted in thick black and the aligned contigs are depicted in thick grey. No contig parts remain unaligned. Pink and yellow boxes indicate the two largest direct repeats Rep1 and Rep2 evincing signature of recombination (thin black lines, some alignments represent an alternative flank composition of Rep1 and Rep2, which breaks the alignment into several parts). The outer contig is constructed based on a single PacBio read, hence it is unlikely to suffer from potential mis-assembly artifacts. b, The mitochondrion assembly graph with protein-coding genes annotated at least partly on one of the two short alternative edges and Rep1 and Rep2 depicted as in (a). Inner orientation of annotated genes reflects the positive strand, while the outer orientation represents the reverse strand genes (black arrows), grouped exons indicate cys-splicing introns opposing to trans-splicing introns between the groups.

The newly assembled mitochondrion genome was annotated based on the previous annotation performed by Petersen et al. (2017). As one major difference, we found that the *cox2* open reading frame is complete rather than truncated as in the previous annotation. It is notable that the *cox2* gene is located across the transition from the common part for both chromosomes and the unique part of Chr1, a feature not contained in the previous assembly (Figure 1B). Furthermore, our annotation includes eight additional tRNA genes. The unique part of Chr2 encodes two essential genes: *cob* and *nad1-exon4*, while the unique part of Chr1 encodes several essential genes: *nad1 exones2-3, nad2, apt8, nad5 exones4-5, ccmB, mttB* and *cox2*. Notably, *nad1-exon3* and *nad1-exon4* are encoded on unique parts of different chromosomes implying an interchromosomal trans-splicing of *nad1-intron3*, therefore both chromosomes are required for Nad1 expression, such that no chromosome is dominant (Figure 1B). The presence of all genes essential to mitochondria function, along with a set of accessory genes in the newly assembled chromosomes serves as an internal validation for the assembly completeness.

### Recombinationally active repeats manifest in eight chromosomes

The PacBio contig alignments to Chr1 and Chr2 further revealed two perfect direct repeats (Rep1 – 4,845 bp and Rep2 – 3,695 bp) and the distribution of PacBio contigs suggested that they would be recombinationally active (Figure 1A). A recombination event would lead to an alternative sequence composition of the repeat flanking regions. The recombination activity of each repeat pair was calculated as the proportion of PacBio reads coverage of the alternative recombined composition from the total PacBio coverage in the repeat region (Supplementary Figure S1). We found high recombinational activity of Rep1 (53.1%) and Rep2 (48.3%). Additional 81 repeats were characterized by a low recombination frequency (<2%) and the remaining 504 repeats had no recombinational activity. Taken together, products of homologous recombination via Rep1 and Rep2 form a pool of main genome chromosomes (Figure 2), whilst 81 repeats with low recombinational activity correspond to additional substoichiometric forms.

**Fig. 2:**
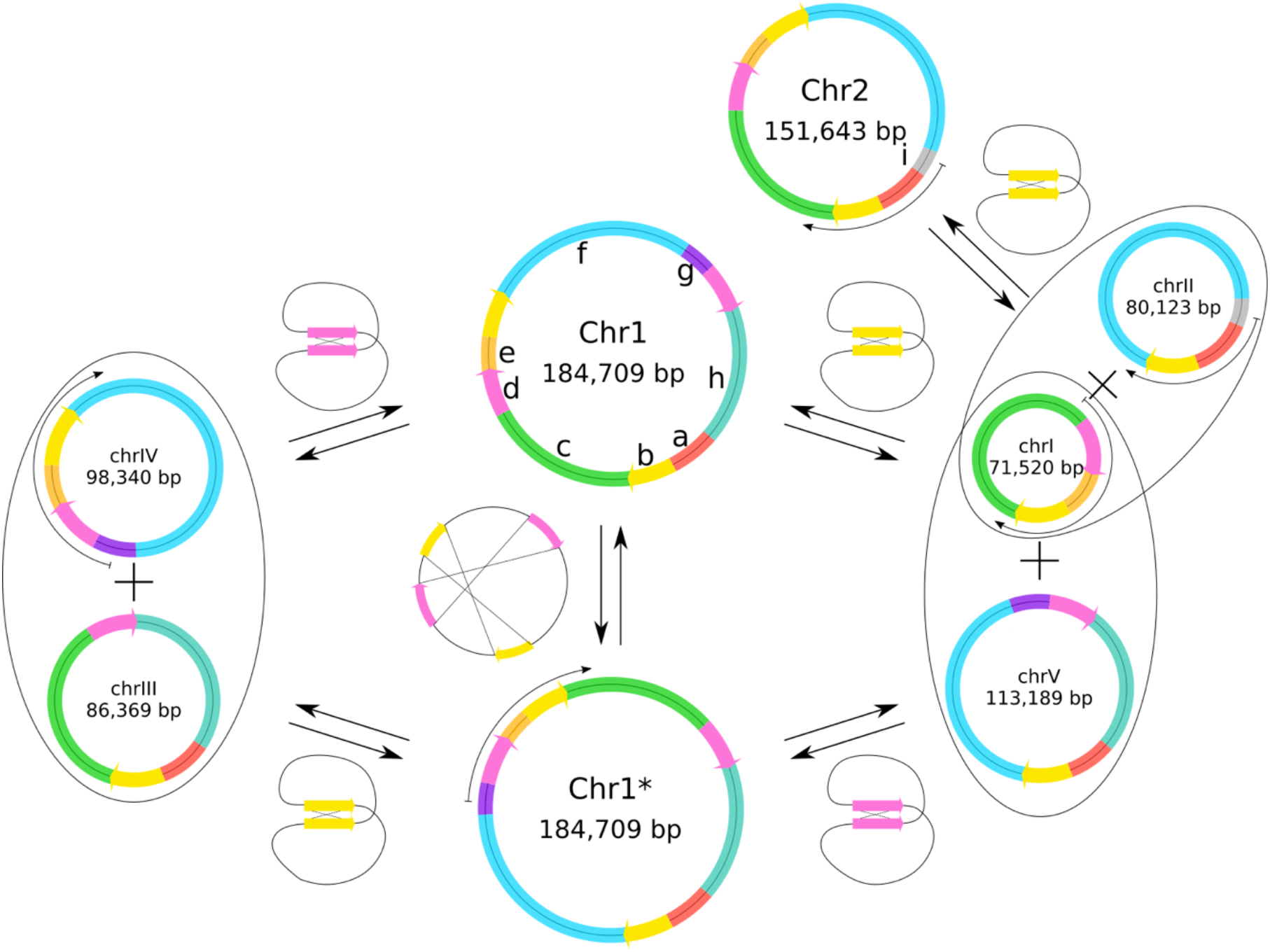
Predicted rearrangements of main genome chromosomes via homologous recombination. Three master circles (Chr1, Chr2, and Chr1*) and five subgenomes (chrI-V) are depicted as circles combined by intact segments, designated by letters a to i, and coloured correspondingly. Segment length ranges between 2,339 and 70,665 bp (see Supplementary Table S1 for details). Thick black arrows mark transformations resulting from reversible recombination reactions via either of the recombinationally active repeats Rep1 (pink arrow) and Rep2 (yellow arrow). Thin black arrows demonstrate examples of PacBio read alignments that confirm the corresponding configuration. All possible configurations are composed of a subset of the five subgenomes. We note that the presence of larger circles resulting from alternative combinations of the described subgenomes cannot be ruled out.

The presence of the recombinationally active repeats indicates the existence of several different chromosomal configurations. Note that recombination of a direct repeat pair splits a master circle into two circular subgenomes (Figure 2). Let us consider Chromosome 1 (Chr1): a recombination via Rep1 leads to subgenomes chrIII and chrIV; similarly, a recombination via Rep2 produces subgenomes chrI and chrV. A recombination via both Rep1 and Rep2, in any order, results in a rearranged conformation of Chr1, termed Chr1*. Chromosome 2 (Chr2) has only Rep2 and therefore only one recombination pathway producing subgenomes chrI and chrII (Figure 2). In total, we predict the main genome to be composed of five subgenomes (chrI-V) and three master circles (Chr1, Chr2, and Chr1*).

To further validate the presence of all eight predicted configurations we used PacBio long read sequencing. For that purpose, all mitochondrial chromosomes were broken into nine genome segments which remain complete in all configurations. Each of the putative chromosomes is thus characterized by a certain order of genome segments **a…i**. PacBio read alignments that cover completely the assigned combination supply a confirmation for the existence of the corresponding chromosome (i.e., segment order). For example, **g-d-e-b-c** segment order appears only in Chr1*, while **c-d-e-b-c** univocally defines chrI (Figure 2). Using this approach, we detected at least one sustaining read for five out of the eight main genome chromosomes: Chr1*, Chr2, chrI, chrII, chrIV (Figure 2, Supplementary Table S2). Notwithstanding, the expected number of covering reads depends on the segment combination total length. The remaining three configurations require considerably longer reads than those available in our data. For the same reason, Chr2 and chrII were confirmed with 97 and 93 reads respectively, whilst Chr1*, chrI, and chrIV were confirmed each by 10 reads or less. The number of supporting reads points towards an equilibrium with equal amount of Chr2 and chrII for the corresponding reversable reaction in the reference sample (Figure 2). Our analysis thus confirms that the *Z. marina* mitogenome contains at least five out of eight predicted chromosomes, nonetheless, their relative abundance and recombination dynamics remain elusive.

### Worldwide population data reveals no evidence for (sub)stoichiometric shifting in the eelgrass mitogenome

The occurrence of (sub)stoichiometric shifting is expected to change the relative abundance of mitochondrial chromosomes over time. To investigate the evolution of chromosome stoichiometry at the population level, we compared the relative mitochondrial chromosome copy number among worldwide *Z. marina* populations. The analyzed data comprises genome sequences from 163 individual *Z. marina* clones sampled in 16 locations in the Pacific and Atlantic oceans (Yu et al. 2022). The sampled tissues comprised early developmental stages, i.e., the meristematic region and the base of first leaflets. To estimate the chromosome copy number, we aligned short-read data from the populations sequencing to the reference mitochondrion genome (as presented in Figure 1) and calculated the absolute copy number of the nine genome segments (**a-i**). Note that the segment copy number is unaltered by the occurrence of Rep1 or Rep2 recombination (Figure 2). The result revealed a similar pattern of the segment copy number within and across populations (Figure 3A). The *Z. marina* phylogeography indicates deep divergence over 350 kya between the Californian populations and the remaining populations; another deep split between the Atlantic and Pacific populations is estimated over 200 kya (Yu et al. 2022; see Figure 3B). The similarity of segment copy number is thus observed across closely and distantly related populations alike. Note that exceptions to the general pattern correspond to samples having a low copy number of each segment (e.g., Alaska-Izembek (ALI) and Alaska-Safety Lagoon (ASL)). One possible trace of (sub)stoichiometric shifting may observed in the Washington state (WAS) population. However, closer inspection of WAS sequence data (12 samples) revealed a segmental duplication in the mitogenome that altered the relative copy number of segments **a** and **h** (Supplementary Figure S2). To examine the conservation of chromosome stoichiometry, we compared the relative copy number of the segments within and among the sampled populations. Our results show that the relative copy numbers of the nine genome segments are homogeneous across all 16 populations (Supplementary Figure S3). Taken together, the similarity in segment copy number across populations suggests an unprecedent conservation of chromosome stoichiometry in *Z. marina* populations worldwide.

**Fig. 3:**
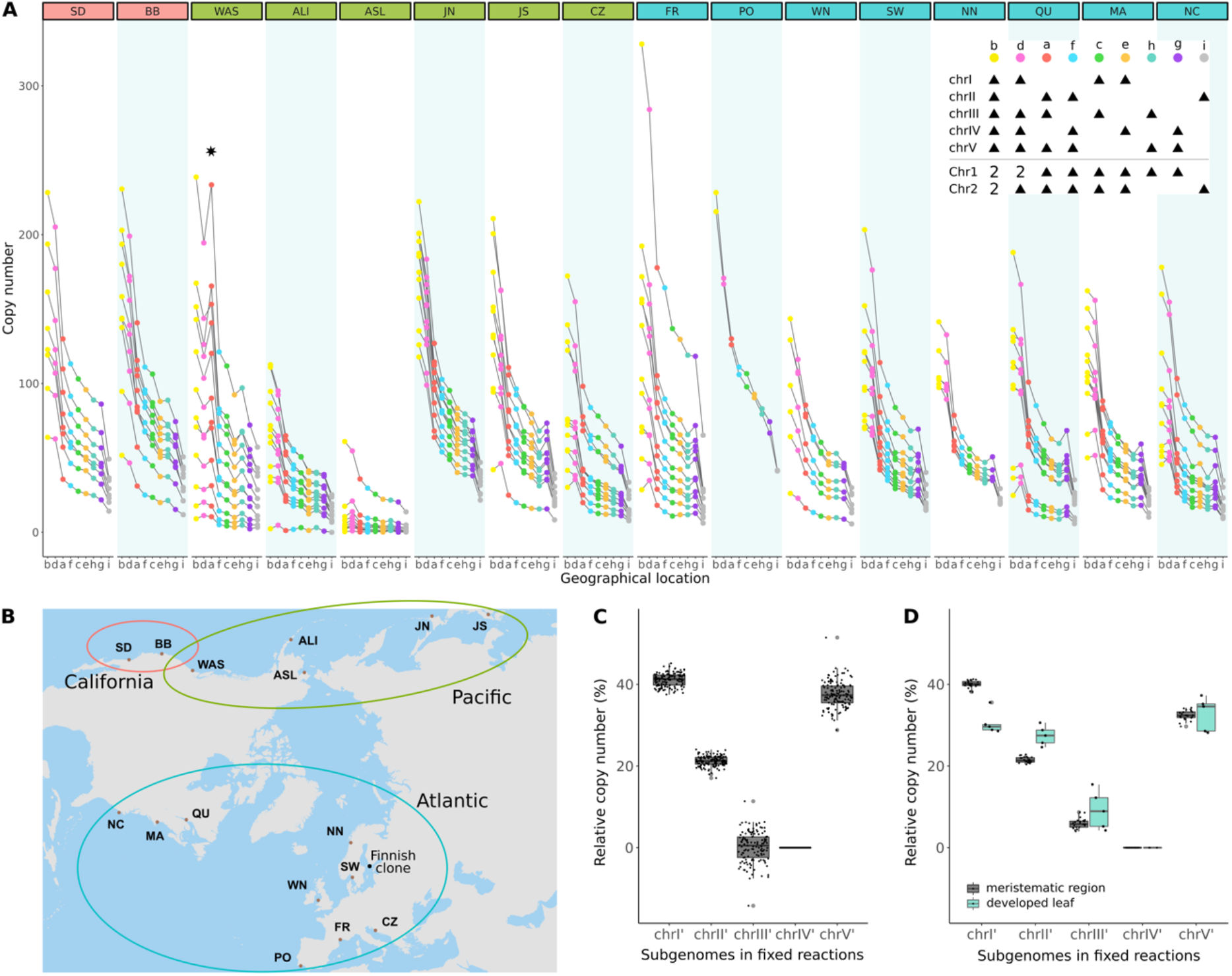
Mitochondrial chromosome stoichiometry is conserved worldwide. a, Absolute copy number of genome segments **a…i** of individual samples grouped by location of sampling. Segment copy number is depicted by dots (colored according to Figure 2); the dots are connected by grey lines within individual samples. An increased segment copy number due to duplication in the WAS samples is highlighted by a star. Segment composition of the chromosomes is shown in the legend (top right) where black triangles show the presence of segments in the corresponding chromosome. Chr1* has identical genome segment content to Chr1. b, Geography of all sampling locations (brown dots) grouped according to the main clades in *Z. marina* phylogeny (Yu et al. 2022). Abbreviations: San Diego, California (SD); Bodega Bay, California (BB); Washington state (WAS); Alaska-Izembek (ALI); Alaska-Safety Lagoon (ASL); Japan-North (JN); Japan-South (JS); North Carolina (NC); Massachusetts (MA); Quebec (QU); Northern Norway (NN); Sweden (SW); Wales North (WN); Portugal (PO); Mediterranean France (FR); Croatia (CZ). Additional black dot indicates the location of Finnish clone in the Archipelago Sea, southern Finland, used for meristems to leaves comparison. c, Relative copy number of the subgenomes chrI’-V’ after fixing all reversible recombination reactions to the absence of Chr1, Chr2, Chr1*, and chrIV calculated for the 132 reliable samples. d, Relative copy number of the subgenomes chrI’-V’ calculated for samples of meristematic region (n=24) and developed leaves (n=5) collected from the Finnish clone (Yu et al. 2020). The calculation was performed after fixing all reversible recombination reactions in the same way as in (c).

### The mitochondrial chromosome stoichiometry suggests equal copy number of the five subgenomes

The conserved relative copy number of genome segments reported here, indicates a stable stoichiometry of the *Z. marina* mitochondrial chromosomes. Note that inferences on chromosome stoichiometry via short read mapping and without additional long read data requires that reversible recombination reactions are considered fixed. For that purpose, we fix the recombination reaction such that the master circles Chr1 and Chr2 are the remaining chromosomes. Using that assumption, the copy numbers of segment ***a*** and segment ***e*** are equal, since both segments are present once in each master circle. However, our data clearly suggests that segment ***a*** has a higher abundance comparing to segment ***e***. Moreover, the relative copy number of segments ***a*** and ***e*** seem to match well their presence in the subgenomes. Segment ***a*** is present in three of the subgenomes and segment ***e*** in only two subgenomes (Figure 2; Figure 3A; Supplementary Figure S3). Notably, the relative copy number of the remaining segments precisely reflect their expected quantity according to the subgenomes segment composition (Supplementary Figure S3). Consequently, we suggest to consider the five subgenomes chrI-V as primary chromosomes in the eelgrass mitogenome. Conversely, the master circles Chr1, Chr2, and Chr1* should be considered as occasional combination of certain subgenomes. Therefore, our assessment of the chromosome stoichiometry focuses only on the five subgenomes chrI-V (see Figure 2).

We calculated the relative copy number of the five subgenomes from the copy number of genome segments **a** to **i**. A direct assessment of the relative abundance of all five chromosomes is impossible due to the possible transformation of chrIII and chrIV to chrI and chrV (Figure 2). For the purpose of our calculation here, we therefore set chrIV copy number to zero. Note that the copy numbers of segments **e, i**, and **g** correspond to the copy number of chrI’, chrII’, and chrV’ respectively. Hence, the copy number of chrIII’ can be measured as the difference in copy number of the three segment pair combinations: **h** and **g, c** and **e**, or **a** and **f** (see Figure 3A legend; Supplementary Figure S4). Estimating chrIII’ copy number using the first combination, **h** and **g**, revealed that the proportion of the five subgenomes chrI’:chrII’:chrIII’:chrIV’:chrV’ slightly deviates from a ratio 2:1:0:0:2 (Figure 3C). Note that under the above assumption, the fraction of chrIV’ is 0, since the copy number of subgenome chrIV had to be set to zero for all samples. The narrow distributions of the subgenome relative abundance among the samples supplies a confirmation for the conservation of chromosome stoichiometry across all populations. If we suppose equal number of pairs chrI-chrV and chrIII-chrIV, we obtain a stoichiometry with a ratio of 1:1:1:1:1 (Supplementary Figure S5). Similar observations of roughly even ratios of the mitochondrial chromosomes have been made for *Silene noctiflora, Silene conica, Psilotum nudum*, and *Lophophytum mirabile* (Sloan et al. 2012; Guo et al. 2017: 20; Sanchez-Puerta et al. 2017). The equality of copy number among the five subgenome in *Z. marina* would correspond to a naturally stable equilibrium.

Repeating the estimation of chrIII’ copy number using the two other segment combinations **c** and **e**, and **a** and **f**, results in a significantly higher relative copy number (Supplementary Figure S6), contrary to expectations based on the segment composition of the main genome chromosomes (Figure 3A). Even if we have to suggest an unknown component of the chromosome composition, the narrow distribution of the estimated chrIII’ relative copy number using above two combinations suggests that the source of deviation is common to all sampled populations. We conclude that the *Z. marina* mitogenome likely contains additional forms that are not composed of the five subgenomes chrI-V; the existence of such alternative forms would alter the relative abundance of certain genome segments.

### Mitochondrial chromosome composition in developed leaves differs from the meristematic region

The striking conservation of mitogenome chromosomes may point towards either a specific property of *Z. marina* due to its compact genome, or a specific property of meristematic region due to its contribution to plant germline. In order to resolve the ambiguity, we analysed additional samples collected from a single *Z. marina* clone from the Archipelago Sea, southern Finland (Yu et al. 2020). The chromosome relative copy number was calculated for 24 samples of the meristematic region and 5 samples of developed leaf tissue, in accordance with the segment composition of the main genome (Figure 3A). The estimated relative chromosome copy number of the five chromosomes in the meristematic samples was within the range of the values calculated from the worldwide population dataset (Figure 3CD). Hence, the mitogenome composition observed worldwide is conserved in the Finnish clone as well. In contrast, the estimated relative copy numbers in developed leaves were markedly different from those observed in the meristematic region, especially for chrI’ and chrII’ (Figure 3D). Furthermore, the leaf samples show a wider distribution of the estimated chromosome copy number, even within a single clone, indicating the occurrence of independent (sub)stoichiometric shifting events during ontogenesis. Thus, the chromosome composition is more diverse between different plant tissues of the same clone than between meristems of populations diverged ca. 350,000 years ago.

## Discussion

Variation in chromosome stoichiometry is a key characteristic of plant mitochondrial genomes where chromosome copy number in mitochondria may alternate by (sub)stoichiometric shifting. A long-standing question is therefore whether and how (sub)stoichiometric shifting translates to the next generation thus contributing to the evolution of mitochondrial genetic diversity over time. Here we demonstrate that the mitochondrial chromosome copy number in *Z. marina* meristematic region remains homogeneous among widely divergent populations, implying a conservation of chromosome stoichiometry in the plant germline over 350,000 years since the last common ancestor of extant populations (Yu et al. 2022). Our results thus reveal a novel perspective on plant population genetics.

Quantifying the stoichiometry of products from chromosome recombination reaction – that is, master circles and their subgenomes – at equilibrium remains challenging. Long read data enabled us to characterize the pool of mitochondrial chromosomes and further reveal evidence for equal copy number of the master circle Chr2 and its subgenome chrII. The measured recombinational activity of Rep1 and Rep2 is close to 0.5 (Supplementary Figure S1), which furthermore points towards equal amounts of the products in all reversible recombination reactions. Our results thus differ from other studies that reported significant superiority in copy number of subgenomes over the master circles (Alverson et al. 2011; Sloan et al. 2012). The relative copy number of master circles to subgenomes may vary among plant tissues and reaches the prevalence of master circles in the plant meristem, which may be essential for the mitogenome heritability (Arrieta-Montiel et al. 2001; Preuten et al. 2010; Woloszynska 2010; Mower, Sloan, et al. 2012). Our observation raises the question whether the stoichiometry of recombination reaction equilibrium shifts rapidly or rather remains stable due to certain regulatory mechanisms, e.g., nuclear control of homologous recombination (Albert et al. 2003; Arrieta-Montiel et al. 2009). Further studies of mitogenome composition using long-read sequencing at the population level are needed in order to assess whether the observed mitogenome conservation is a unique feature for the eelgrass, or rather a general phenomenon.

The conventional way for describing a mitochondrial genome is to present master circles as primary chromosomes and subgenomes as occasional products of homologous recombination, even if the subgenomes eventually dominate in quantity. This approach incorporates an assumption that subgenomes can be grouped into pairs to form the master circles. However, our data supports the opposite perspective: subgenomes, rather than master circles, should be considered as the relevant genetic entities, similar to observations in the monkeyflower mitogenome (Mower et al. 2012). Yet, our analysis of segment copy number suggests that focusing on subgenome may still supply an incomplete mitogenome representation. The transformation of genome segment copy number into the chromosome stoichiometry revealed an unexpected significant difference using different formulations (Supplementary Figure S6), all based on possible rearrangements of the main genome chromosomes (Figure 2). This result suggests the presence of additional configurations that affect the genome segment abundance but are not represented in the main genome (i.e., these are being generally underestimated). The conventional depiction of plant mitogenomes solely as master circles and subgenomes thus supplies a partial representation of the chromosome composition diversity.

To achieve a complete mitogenome representation we suggest two alternative hypotheses for the source of deviation in segment copy number from the expectation according to the repertoire of main genome chromosomes. The first possible explanation to our conundrum is an effect of substoichiometric forms. Although each substoichiometric molecule persists in a much lower proportion than the main genome chromosomes, the substoichiometric reservoir altogether could be responsible for the shifts in genome segment relative copy numbers. For example, an underrepresentation of segment **f** in substoichiometric forms, would lead to an overestimation of chrIII’ copy number when using the **a**-**f** formulation (Figure 3D). We note, however, that since the variation of **a**-**f** among populations is rather small, this explanation implies that the composition of the substoichiometric forms reservoir remains stable across all populations, in agreement with previous suggestions in the literature (Laser et al. 1997; Arrieta-Montiel et al. 2001; Woloszynska and Trojanowski 2009; reviewed in Woloszynska 2010). An alternative possibility is the presence of partially overlapping subgenomic linear molecules instead of, or in addition to, complete circular molecules (reviewed in Gualberto and Newton 2017; Arimura 2018). Overrepresented genome segments may thus correspond to overlapping regions of subgenomic linear molecules. For example, if such overlap of linear forms is found within segment **a**, then this will lead to an overestimation of segment **a** copy number and therefore a deviation of **a**-**f** from our expectation. Considering that linear molecule ends are proposed to be recombination-dependent replication start points, the small variation in **a**-**f** under this scenario, suggests that the origins of replication are conserved throughout all populations.

The mechanisms controlling mitochondrion chromosome stoichiometry remain unclear and are most likely a combination of several factors. Two mechanisms have been discussed in the literature: (i) the effect of nuclear control of recombination reaction dynamics (Albert et al. 2003; Arrieta-Montiel et al. 2009) and (ii) having chromosome conformations that are specific to meristematic tissues, which may reduce the possibility of (sub)stoichiometric shifting (Arrieta-Montiel et al. 2001; Woloszynska 2010; Liberatore et al. 2016). Additional possibility is section pressure related to change of gene dosage. Relative expression of genes located on chromosomes having different abundance might have a gene dosage effect, which could be subject to stabilizing selection, similarly to observations made for prokaryotic plasmids (Nicoloff et al. 2019).

Our study reveals a striking conservation of the mitochondrial chromosome composition and stoichiometry in meristematic regions across worldwide populations. This observation is all the more surprising given the complex chromosome architecture revealed here. At the same time, developed leaf samples exhibit evidence for (sub)stoichiometric shifting in comparison to the meristematic regions of the same clone, in agreement with previous observations in *Phaseolus vulgaris* (Woloszynska et al. 2012). Given the above we conclude that *Z. marina* germline is preserved from (sub)stoichiometric changes and therefore they are not transmitted to next generations neither vegetatively nor by sexual reproduction. However, the mitochondria chromosome composition and gene order are highly variable among plant species, even those that are closely related (Palmer and Herbon 1988; Mower, Sloan, et al. 2012). Consequently, we suggest that changes in chromosome stoichiometry are not a major driver in the evolution of mitochondrial chromosome architecture in eelgrass. Instead, we predict intrachromosomal rearrangements, e.g., duplication, translocation, and inversions, as main events for mitogenome evolution during Zosteraceae species divergence, as observed in the WAS samples. Notably, rearrangements of the mitochondrial chromosomes in *Brassica napus* lines may lead to a cytoplasmic male sterility (Chen et al. 2011), thus supporting a role of such rearrangements in speciation events. Substoichiometric shifting, in contrast, may be a plastic property of mitochondrial genomes in somatic tissues that is inhibited in the germline by specific buffering mechanisms. Our findings open up avenues for further research on the evolution of chromosome architecture and stoichiometry in plant mitochondria, its maintenance, as well as possible implications for plant diversification.

## Materials & Methods

### Data source

The chromosome-level nuclear and chloroplast *Z. marina* genome assembly was downloaded through the ORCAE platform (https://bioinformatics.psb.ugent.be/gdb/zostera/) and the long-read PacBio sequencing data were downloaded from NCBI, accession number: PRJNA701932 (Ma et al. 2021). The Illumina whole-genome sequencing of *Z. marina* meristematic region of the inner leaf base was obtained from JGI, Proposal ID: 503251 for the worldwide population dataset and from NCBI (BioProject: PRJNA557092) for the Finnish clone dataset (Yu et al. 2020; Yu et al. 2022). Worldwide population dataset samples were preselected according to (Yu et al. 2022) to avoid representatives of the same clone and selfing. In addition to the published data sets of the global population-level analysis, and the 24 clone mates at Ängsö, additional plants of the Finnish clone were collected in 40×40 cm areas in summer 2021 by SCUBA diving along a horizontal transect line in 2m water depth at locations ca. 20 m distant from one another. Approximately 80 mg of dried leaf tissue (deliberately excluding the meristematic reagion) from a total of 6-8 leaf shoots at each spot were extracted as in (Yu et al. 2022) and subjected to full genome resequencing using a paired end PE150 library preparation and an Illumina HiSeq X Ten sequencing platform to a coverage of approximately 50x at BGI Hongkong. Sequences have been submitted to NCBI Short Read Archive under accession no xxxx).

### Mitochondrial genome assembly

PacBio long reads were assembled into contigs by Canu with default parameters and expected genome size of 230Mbp (Koren et al. 2017). The result contigs were aligned to the mitogenome assembly from Petersen et al. 2017 (RefSeq: NC_035345.1) by nucleotide BLAST (version 2.2.28) with default parameters (Altschul et al. 1990). Contigs with the over 10 Kbp long alignment to the subject mitogenome, including nine contigs with the length range of 49,378-115,620 bp, were considered mitochondria-related and used for further assembly. The assembly process was based on manual overlap of the nine contigs resulted in the two-edge assembly graph (Figure 1). Self-duplications and breaks in several alignments are found next to the Rep1 and Rep2 repeats; self-duplications are likely to be the result of a contig misassembly, whilst the breaks may supply evidence for alternative configurations. All nine contigs could be aligned to the proposed mitochondrial genome assembly graph entirely, thereby excluding any additional ambiguities (Figure 1). The mitogenome circularity was confirmed by discontinuous coverage by PacBio contigs (Figure 1) and Illumina short reads (as suggested in Petersen et al. 2017). Moreover, the mitochondrial genome proposed in Petersen et al. is included in the new assembly, rearranged and slightly extended. All together the assembly graph on Figure 1 is likely to represent complete *Zostera marina* mitogenome with the summary length of the edges of 187,048 bp resulting in a considerably compact but within the range of plant mitochondrion genomes.

Further validation of the mitogenome sequence was performed by Illumina reference short reads, which were obtained from Olsen et al. 2016 (SRR3926352). For each selected PacBio contig Illumina reads were used to confirm absence of relative coverage drops which would point at a chimeric origin of the assembly. Low-quality contig ends were omitted from the final genome reconstruction. Coverage was calculated directly by SAMtools from the raw read mapping performed by BWA-MEM with default parameters (Li 2013; Danecek et al. 2021). To overcome edge effects on coverage, Illumina reads were aligned to two different linear forms of the joined mitochondria assembly with non-repeated break points of the circles. Then the final coverage was calculated per position as maximum coverage of the corresponding position in two shuffles. For the proposed assembled mitogenome a connection heat map was created: the genome was subdivided into blocks of 100 bp, then for each pair of blocks the number of read pairs aligned to both blocks simultaneously was counted. Both coverage and connections were used as additional confirmation of correctness and completeness of the mitochondrion assembly.

### PacBio reads for chromosome validation

Recombination activity of repeats was quantified with the use of PacBio long reads. Potential recombination sites were identified by aligning the assembled mitochondrion to itself by BLAST with the minimum repeat length of 50 bp, minimum identity 80% and minimum distance between the two copies of 2000 bp A total of 380 direct and 245 inverted repeat pairs were identified. A recombination database was constructed, containing four possible combination of 1000 bp flanks for each repeat pair: M1, M2 – flank composition as they are in the assembly master circles; R1, R2 – recombined versions. All PacBio reads were aligned to the recombination database by BLAST with the minimum identity 80% and minimum flank coverage 80%. The proportion of reads aligned to R1 or R2 was used as an estimate of the recombination activity of a particular repeat pair. Because of a certain region complexity, we additionally marked repeats, which are nested in Rep1 or Rep2 and therefore might show a misleading recombination signal. For the majority of repeat pairs, the proportion of recombined configurations was below 2% but above zero, hence, corresponding PacBio reads are likely to originate from substoichionmetric forms.

Another approach engaging PacBio long reads for chromosome validation was a direct alignment of reads to all main genome chromosomes with a further filtration of unambiguously defined read-chromosome pairs. Such pairs occur when a PacBio reads covers a unique sequence found in only one main genome chromosome. The minimum length of a unique sequence to confirm a chromosome vary significantly, therefore the numbers of sustaining reads are comparable only among Chr1*-chrI -chrIV group (24,730 bp minimum length required) and between Chr2 and chrII (7,121 bp minimum length required). Corresponding numbers of PacBio reads confirming comparable chromosomes reflect their relative abundance in the reference sample. For Chr1, chrIII, and chrV the minimum length of a PacBio read to cover an unambiguous chromosome region to be properly paired is over 39,579 kbp, which dramatically decreases the probability to find at least one suitable PacBio read, explaining the absence of such reads in our dataset.

### Mitogenome annotation

Mitogenome annotation was performed by tBLASTn of protein sequences of other known mitogenomes within order Alismatales including *Z. marina* previous assembly (Petersen et al. 2017), tRNAs were additionally annotated by Mitofy (tRNAscan-SE) (Lowe and Eddy 1997; Alverson et al. 2010) and ARAGORN (Laslett 2004). In comparison with the previous assembly the annotation of *cox2* gene appeared as a complete ORF and 8 additional tRNA genes were identified. DNA regions shared with nuclear chromosomes (NUMTs) and chloroplasts (mtptDNA) were detected by BLAST with minimum identity 85% and maximum e-value 1e-10.

### Worldwide population dataset and Finnish clone dataset analysis

Data of Illumina sequencing of 163 samples from 16 populations was retrieved from a previous study of *Z. marina* populations (Yu et al. 2022). Raw reads were mapped by BWA-MEM with default parameters to 67 core nuclear single copy genes longer than 3000bp to calculate target coverage of the sequencing per single replicon which ranges from x2.5 to x55.5. Similarly, raw reads were aligned to the assembly mitogenome graph genome content. Genome segments were defined as unbreakable parts of the mitochondrion – 9 in total counting Rep1 and Rep2 only once (Figure 2). Genome segment copy numbers were assessed as median coverage of non-NUMT and non-mtptDNA positions normalized on one replicon target coverage of the corresponding sample. For the chromosome stoichiometry calculation and the sample pair analysis only non-WAS samples with the minimum coverage of segment **b** (Rep2) of 40x were selected, including 132 samples in total. Finnish clone meristematic region samples (n=24) and developed leaf samples (n=5) were analyzed in the same way as the worldwide population dataset.

## Acknowledgements

We thank Giddy Landan, Ana Garoña, Nils Hülter and Lei Yu for fruitful discussions and critical comments on the manuscript. Sarah Rühmkorff and Christian Pansch-Hattich helped with sampling additional leaf tissue of the Finnish clone at Ängsö. Lei Yu helped with acquiring the short read data from leaf samples. The first author is funded through the Helmholtz School for Marine Data Science (MarDATA), Grant No. HIDSS-0005. The original DNA sequencing was funded by the U.S. Department of Energy (DOE) Joint Genome Institute (JGI) Community Sequencing Program (CSP 502951, 2016, Population and evolutionary genomics of host-microbiome interactions in *Zostera marina* and other seagrasses). The work (proposal: 10.46936/10.25585/60000773) conducted by the U.S. Department of Energy Joint Genome Institute (https://ror.org/04xm1d337), a DOE Office of Science User Facility, is supported by the Office of Science of the U.S. Department of Energy operated under Contract No. DE-AC02-05CH11231.

## Author contributions

MK, TBHR and TD conceived the study. MK designed the approach and performed the data analysis. All authors interpreted the results and wrote the manuscript.

