## Supplementary materials for "Worldwide population genomics reveal long-term stability of the mitochondrial chromosome composition in a keystone marine plant"

### Supplementary information

#### Supplementary Tables

**Supplementary Table S1: Genome segment lengths. Genome segments** are defined as unbreakable genome sequences in main genome chromosomes (Main Fig. 2); **segment color on Fig2** is the visual representation of corresponding segment on Main Fig. 2; **segment length (bp)** is the corresponding total length of the sequence; **non-shared segment length (bp)** is the number of non-NUMT and non-mtpt positions which were incorporated in the copy number calculation. *Zostera marina* is characterized by a small mitochondrial genome as predicted in Petersen et al. 2017 similar to the compact nuclear genome (Olsen et al. 2016; Ma et al. 2021). However, a uniform way of calculating the mitogenome size in case of a multipartite architecture is not settled. A simple length summary of the primary chromosomes chrI-V yields a total size of 449,541bp, however using this formulation, specific genome segments are counted multiple times. If calculated as total length of the master circles only, excluding the subgenomes or alternative rearrangements (Alverson et al. 2011), the total mitochondrial genome size is 336,352bp, which is the length of Chr1 and Chr2 together (that similarly includes repeated genome content). A better representation of the mitogenome size should be the total length of the optimal assembly graph (Varré et al. 2019), namely 187,048bp, or in the extreme case, 178,508bp, if each genome segment **a..i** is counted once.

| Genome segment | Segment color on Fig2 | Segment length (bp) | Non-shared segment length (bp) |
|---|---|---|---|
| **a** | red | 3,424 | 3,309 |
| **b** | yellow | 3,695 | 2,844 |
| **c** | green | 46,792 | 37,493 |
| **d** | pink | 4,845 | 3,102 |
| **e** | sand | 16,188 | 8,160 |
| **f** | blue | 70,665 | 31,845 |
| **g** | violet | 2,947 | 1,430 |
| **h** | marine | 27,613 | 25,030 |
| **I** | grey | 2,339 | 2,174 |

**Supplementary Table S2: Long read alignments for chromosome confirmation. Chromosomes** are the eight main genome chromosomes; **unique genome segment combination** is the minimum in length order of genome segments represented exclusively in the corresponding chromosome; **minimum read length (bp)** is the minimum long read length to confirm the corresponding chromosome; **number of aligned PacBio reads** is the number of reads which confirmed the corresponding chromosome being aligned to the identified unique genome segment combination.

| Chromosome | Unique genome segment combination | Minimum read length (bp) | Number of aligned PacBio reads |
|---|---|---|---|
| Chr1 | **g-d-h-a-b-c** | 39,579 | 0 |
| Chr2 | **i-a-b-c** | 7,121 | 97 |
| Chr1* | **g-d-e-b-c** | 24,730 | 10 |
| chrI | **c-d-e-b-c** | 24,730 | 5 |
| chrII | **i-a-b-f** | 7,121 | 93 |
| chrIII | **c-d-h-a-b-c** | 39,579 | 0 |
| chrIV | **g-d-e-b-f** | 24,730 | 3 |
| chrV | **f-g-d-h-a-b-f** | 42,526 | 0 |

#### Supplementary Figures

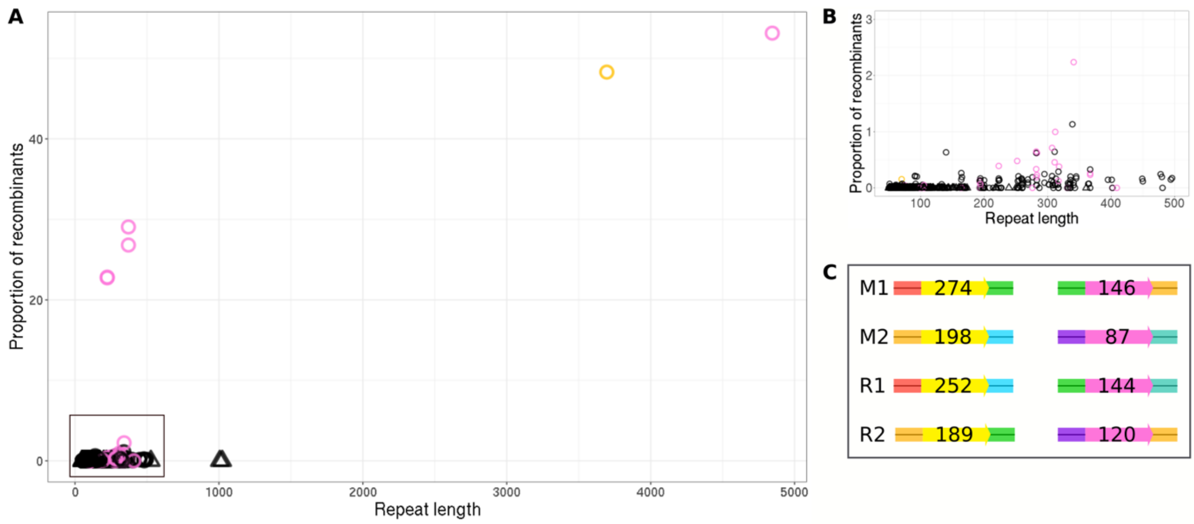

**Supplementary Fig. S1: Recombination activity analysis of all large and intermediate size repeat pairs within Z. marina mitogenome.** a, Recombination activity of all intermediate size and large repeat pairs. Circles indicate direct repeats, triangles – inverted. Repeats nested in (or equal to) Rep1 and Rep2 are colored in pink and yellow correspondingly, such intermediate size repeats might show a signal which is a reflection of the Rep1 and Rep2 recombinational activity and therefore cannot be taken into account. b, A zoom into the bottom-left part of the graph on a, with multiple repeats demonstrating weak recombinational activity and likely referring to the substoiciometric reservoir. c, The master (M1, M2) and alternative (R1, R2) flank compositions of the repeats Rep2 and Rep1 with the numbers of PacBio reads supporting the corresponding flank composition.

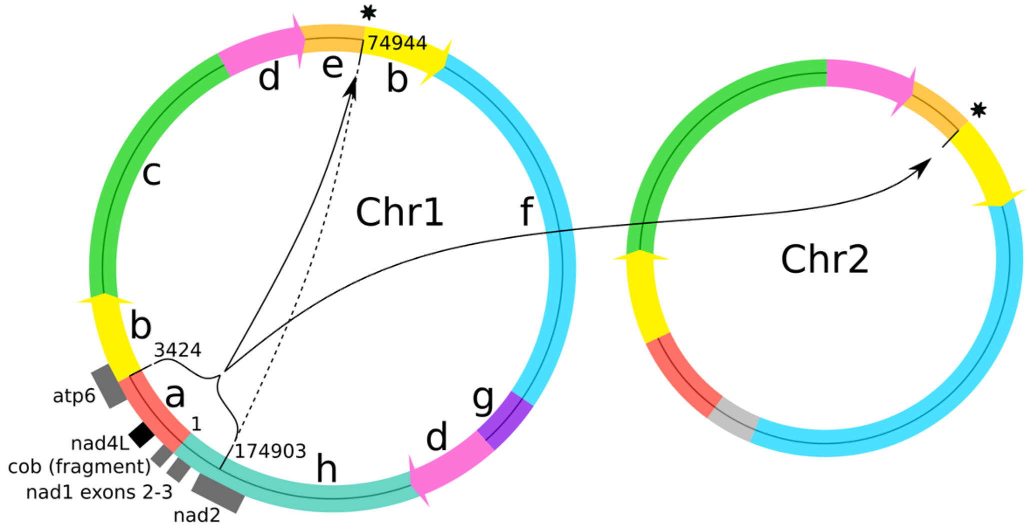

**Supplementary Fig. S2: Intrachromosomal duplication detected in all WAS samples.** Short Illumina reads aligned to the reference genome no linkage between segments **e** and **b** in all WAS samples, therefore pointing towards a breakpoint between **e** and **b** on all chromosomes (marked with a star). Instead, an additional contact was identified between the position 174903 in segment **h** and the beginning of segment **b** (dashed line). Given the doubled coverage of the region 174903-3424 we concluded that this region was duplicated in between segments **e** and **b** on both Chr1 and Chr2 and all their derivatives that have consequent segments **e** and **b** in all other samples (black arrows). Grey rectangles show all genes that partially overlay with the duplicated region. The black rectangle indicates nad4L gene duplicated in a complete form proposing a possible increase of its expression.

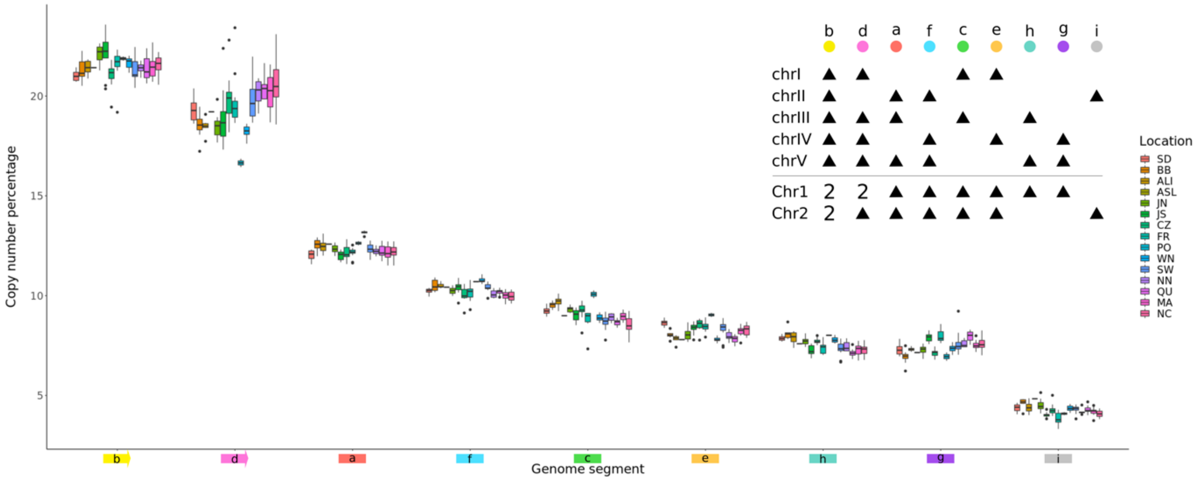

**Supplementary Fig. S3: Relative abundance of genome segments across 15 populations worldwide.** For all non-WAS samples, the relative abundance (Y axis) of the nine genome segments **a…i** (X axis) was grouped by location (colored boxplots) and ordered from California via Pacific to Atlantic from left to right. On the legend in the top-right corner black triangles show the presence of genome segments in the corresponding chromosome, Chr1* has identical genome segment content to Chr1.

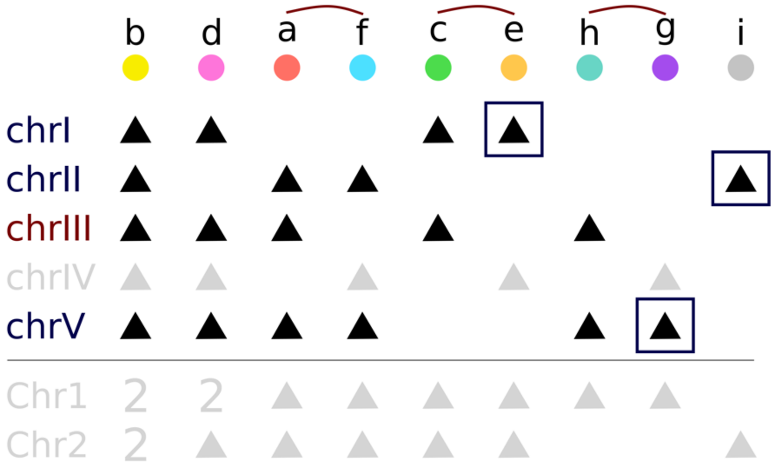

**Supplementary Fig. S4: Scheme of the nine genome segment distribution among the main genome chromosomes when fixing the reversible reactions to zero master circle and chrIV products.** The scheme is similar to the legend on the main Fig. 3A and the Suppl. Fig. 3. The violet squares indicate genome segments univocally representing subgenomes chrI, chrII, and chrV. The brown arcs indicate simple substraction for the chrIII abundance calculation.

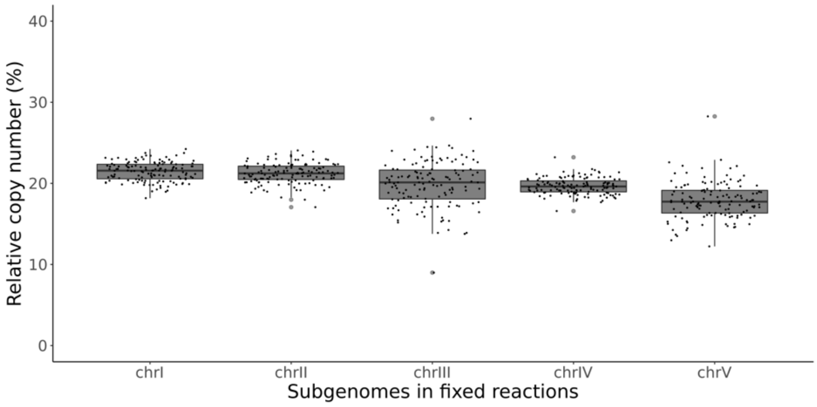

**Supplementary Fig. S5: Relative copy number of the subgenomes chrI-V after fixing all reversible recombination reactions to the absence of Chr1, Chr1*, and Chr2 and exactly half of the chrI-chV pairs being transformed to chrIII-chrIV pairs.** The chrIII abundance calculation is based on h-g method similar to Main Fig. 3C. Only the 132 reliable samples are taken into account.

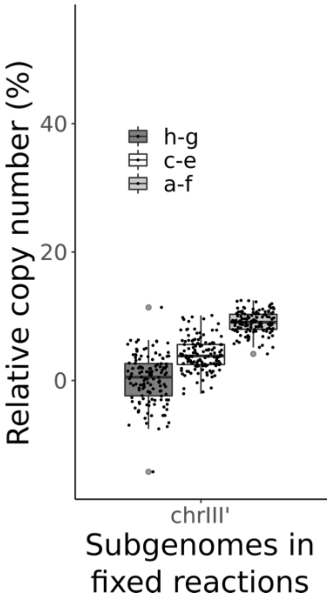

**Supplementary Fig. S6: Relative copy number of the subgenome chrIII after fixing all reversible recombination reactions to the absence of Chr1, Chr1*, Chr2 and chrIV.** The chrIII relative abundance is calculated as **h-g**, **c-e**, and **a-f**. The three methods provide significantly different results with Kruskal-Wallis test p-value below 2.2e-16. Only the 132 reliable samples are taken into account.
